# Imaging of lactate metabolism in retinal Müller cells with a FRET nanosensor

**DOI:** 10.1101/2022.10.05.510984

**Authors:** Víctor Calbiague García, Yiyi Chen, Bárbara Cádiz, François Paquet-Durand, Oliver Schmachtenberg

**Author notes:** Corresponding Author: Dr. Oliver Schmachtenberg, CINV, Instituto de Biología, Facultad de Ciencias, Universidad de Valparaíso, Avda. Gran Bretaña 1111, 2360102 Valparaíso, Chile.

## Abstract

Müller cells, the glial cells of the retina, provide metabolic support for photoreceptors and inner retinal neurons, and have been proposed as source of the significant lactate production of this tissue. To better understand the role of lactate in retinal metabolism, we expressed a lactate and a glucose nanosensor in organotypic mouse retinal explants cultured for 14 days, and used FRET imaging in acute vibratome sections of the explants to study metabolite flux in real time. Pharmacological manipulation with specific monocarboxylate transporter (MCT) inhibitors and immunohistochemistry revealed the functional expression of MCT2 and MCT4 in Müller cells. The introduction of nanosensors to measure key metabolites at the cellular level may contribute to a better understanding of heretofore poorly understood issues in retinal metabolism.

---

Adequate metabolism is a prerequisite for optimal cell function. This issue is critical in tissues with high energy demands, for which the retina is a prime example (Ames et al., 1992; Country, 2017). Interestingly, Warburg’
ss studies from 1924 pointed out that tumor cells and the retina share the use of aerobic glycolysis for energy production, converting almost 70% of consumed glucose into lactate, despite the presence of all elements necessary for the much more efficient oxidative phosphorylation (Wang et al., 1997; Warburg, 1924). However, the reasons for this peculiar metabolism, the specific cell types performing aerobic glycolysis in the retina, and the conditions under which it may occur remain unknown to date. Several studies concluded that Müller cells (MC), the glial cells of the retina, are the source of retinal lactate which is supplied to neurons, especially photoreceptors (Poitry-Yamate et al., 1995; Tsacopoulos et al., 1988). However, a direct measurement of retinal lactate flux in real-time and with cellular resolution has not been reported to date.

Here, we expressed the FRET lactate nanosensor Laconic in organotypic retinal explant cultures to study the lactate dynamics in MCs and compared these with glucose levels assessed with the glucose nanosensor FLII12Pglu-700Δ6, hereafter named Δ6. These types of nanosensors have become popular in recent years for the analysis of metabolic flux and transport of different metabolites in neurons and astrocytes, either in culture or *in vivo* (Mächler et al., 2016; San Martín et al., 2013). In the present study, we applied a functional approach to characterize the expression of different monocarboxylate transporters (MCTs) in MCs. The successful expression of nanosensors in the retina and the results of our analysis may help to further understand the complex, yet enigmatic retinal metabolism at a single cell level, with ramifications for pathological conditions such as diabetic retinopathy.

In this study, healthy 9-day old C57BL/6 mice were used irrespective of sex or weight. Animals were born and raised in the animal facility of the University of Valparaiso, held at 20 – 25°C under a 12h photoperiod with water and food *ad libitum*. The experimental protocols were approved by the bioethics committee of the University of Valparaiso and in accordance with the Chilean animal protection law. To isolate the retinas, mice were deeply anesthetized with isoflurane (Sigma Aldrich) before being sacrificed by decapitation.

Retinal explants obtained from post-natal (p) day 9 wild-type mice were cultured as previously described (Belhadj et al., 2020; Calbiague et al., 2020; Valdés et al., 2016). Briefly, the eyes were treated for 15 min with 0.12% Proteinase K (Cat. No. P2308, Sigma-Aldrich) at 37°C for isolation of the retina together with its retinal pigment epithelium (RPE). Then, the eyes were placed for 5 min in DMEM medium with 10% fetal bovine serum (FBS) to deactivate Proteinase K. The retinas were separated from the choroid and placed with the RPE facing down on cell culture inserts (Millicell, Cat. No. PICM0RG50, Merck Millipore) with DMEM culture medium (Cat. No. 31600034, Thermo Fisher Scientific), containing 10% FBS (Sigma-Aldrich) with 15 mM glucose, which was replaced every two days. The cultures were incubated at 37°C in 5% CO_2_, and 95% humidity for 14 days in a water-jacketed incubator (Thermo Scientific). At p11 and p12 the explants were transduced by overnight incubation with 5×10^6^ plaque-forming units (PFU) of Ad Laconic, AAV-Laconic or Δ6, and imaged after two weeks in cultures. Adenoviral serotype vectors encoding FRET nanosensor Ad FLII12Pglu-700Δ6 (Takanaga et al., 2008)) and Ad Laconic (San Martín et al., 2013) were a gift from Dr. Ivan Ruminot from the Centro de Estudios Científicos (CECs) in Valdivia, Chile. AAV-GFAP-Laconic was constructed by the viral vector facility of ETH Zurich (Laconic: Addgene #44238; hGFAP promoter fragment: DOI: 10.1002/glia.20622).

For retinal slice preparation, the explants were separated from the culture inserts and placed in a chamber with extracellular solution, containing (in mM): 119 NaCl, 23 NaHCO_3_, 1.25 NaH_2_PO_4_, 2.5 KCl, 2.5 CaCl_2_, 1.5 MgSO_4_, 20 glucose and 2 Na^+^ pyruvate, aerated with 95% O_2_ and 5% CO_2_, Ph 7.4. The tissue was embedded in type VII agarose (Cat. No. 39346-81-1 Sigma Aldrich) and cut with a vibratome (Leica VT1000S) to 200 μm thickness. The slices were transferred to the microscope recording chamber, sustained by a U-shaped platinum wire and superfused with oxygenated extracellular solution at room temperature (20°C) under photopic conditions. For live cell imaging, retinal slices were maintained in extracellular solution containing (in mM): 119 NaCl, 23 NaHCO_3_, 1.25 NaH_2_PO_4_, 2.5 KCl, 2.5 CaCl_2_, 1.5 MgSO_4_, 5 glucose and 1 lactate and, aerated with 95% O_2_ and 5% CO_2_, pH 7.4. The lactate concentration was chosen not to saturate the lactate sensor. Since there is only a limited number of commercial MCT inhibitors available, based on prior reports (Murray et al., 2005; Ovens et al., 2010; Vélez et al., 2021), we used three potent and specific inhibitors to isolate MCT isoforms: SR-13800 for MCT1 (Cat. No. 5431, Tocris), AR-C155858 (Cat. No. 4960, Tocris) for MCT1 and MCT2, and Syrosingopine (Cat. No. SML1908, Sigma Aldrich) to inhibit MCT1 and MCT4.

Since the lactate and glucose sensors used here have similar excitation/emission spectra (Laconic: 460/492 nm for mTFP and 515/526 for Venus; Δ6: 440/480 nm for CFP and 513/530 nm for YFP), retinal slices were excited at 430 nm and visualized at 480 nm and 530 nm peak wavelength, as previously reported (Fig. 1A) (Barros et al., 2014; San Martín et al., 2013). All experiments were performed at room temperature (22–25°C) with an upright fluorescence microscope (Olympus BX51) equipped with a 40x water-immersion objective, an Optosplit II emission image splitter (Cairn, UK), and a Sensicam QE digital camera (Cooke Corp.). Data acquisition was performed by custom software written in Python 4.0.1. At the end of the experiments, data were exported for off-line analysis of fluorescence intensities from each emission channel. To obtain the FRET ratio for Laconic (mTFP/Venus) and Δ6 (YFP/CFP), fluorescence intensity values from each ROI and background were measured in ImageJ, version 1.52p (NIH, RRID: SCR_003070). In this study, all ROIs were chosen from the somas of MCs (Fig. 1B and D). The FRET data are displayed as the relative FRET ratio, in percentage of change over time of single experiments.

**Fig 1.**
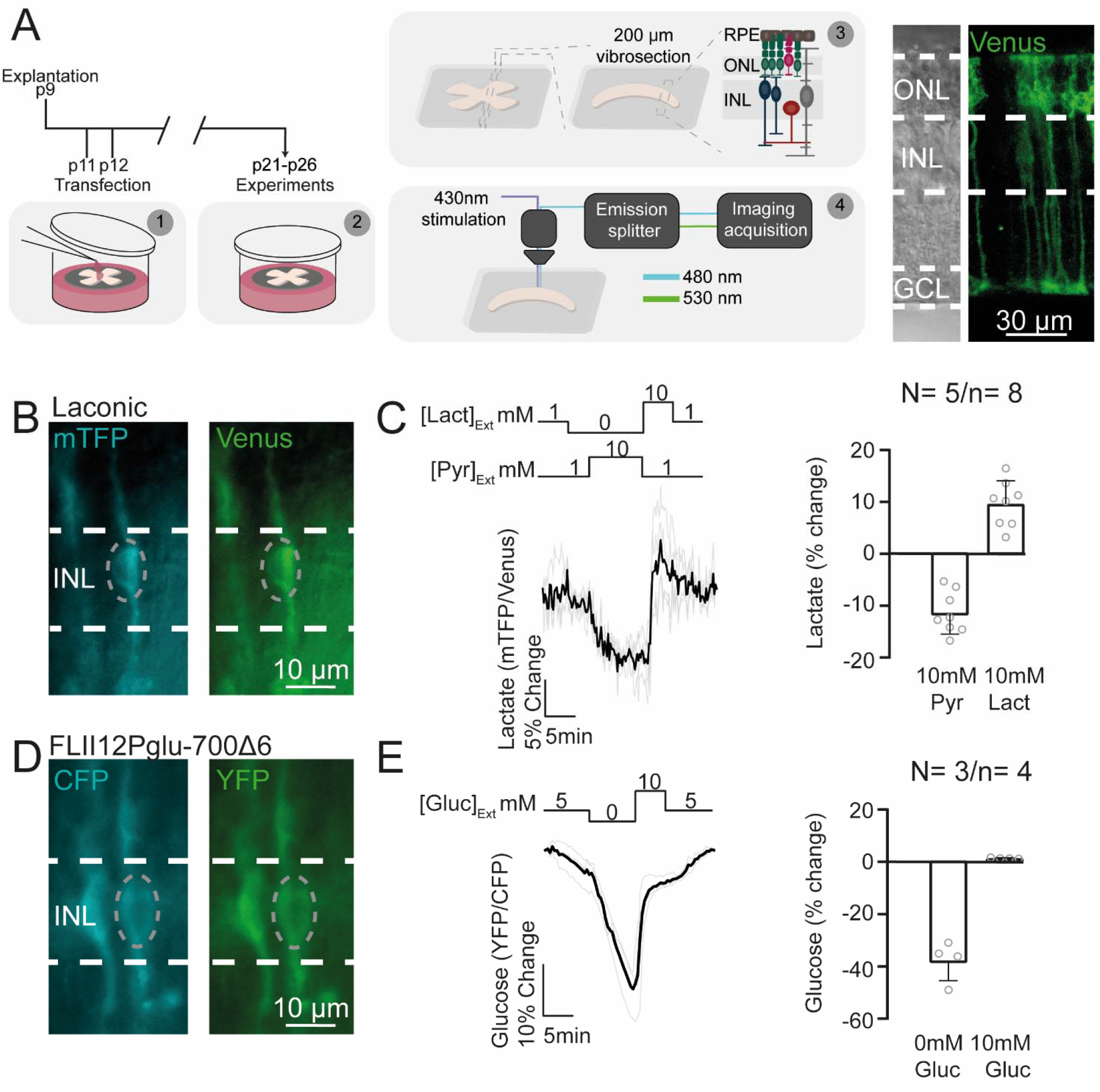
The Laconic nanosensor can be functionally expressed in retinal MCs. (A, left) Scheme of the transfection protocol and the experimental setup. (A, right) Confocal image of Laconic expression in retinal explants after two weeks in culture. (B and D) Representative images of the expression of the lactate sensor (Laconic) and glucose sensors (FLII12Pglu-700Δ6). Dash circles show the area recorded. (C and E) Dynamic range of the lactate and glucose sensor in MCs. The light grey traces represent responses of single cells, and the black trace represents their average. The number of experiments is represented as: N= number of explants; n= number of cells recorded. ONL= outer nuclear layer; INL= inner nuclear layer; GCL= ganglion cell layer; MC(s)= Müller cell(s).

Retinas from p30 mice were fixed in 4% paraformaldehyde for 45 minutes, washed in PBS, cryopreserved with PBS containing 10, 20, and 30% sucrose successively, before embedding in tissue-freezing medium, and stored frozen at -20°C. Transverse sections of 14 µm diameter were obtained with a cryostat (Leica CM-1900) and deposited on poly-L-lysine-coated slides, which were dried at 37°C for 30 min and rehydrated for 10 min in PBS at room temperature (RT). For immunofluorescent labelling, the tissue was incubated in blocking solution (10% normal goat serum, 1% bovine serum albumin in 0.3% PBS-Triton X 100) for 1 h at RT. The primary antibodies: Anti-MCT1 (SLC16A1) (Cat. No. AMT-011, Alomone labs, RRID: AB_2756669), Anti-MCT2 (SLC16A7) (Cat. No. AMT-012, Alomone labs, RRID: AB_2340997), Anti-MCT4 (SLC16A4) (Cat. No. AB180699, Abcam), Anti-Glutamine synthetase (Cat. No. AB73593, Abcam, RRID: AB_2247588) and Anti-glutamate-aspartate transporter (Cat. No. AGC-021, Alomone labs, RRID: AB_2039885) were diluted 1:100 in blocking solution and incubated at 4°C overnight. The slides were washed 3 times for 10 min each with PBS. Subsequently, the secondary antibody, diluted 1:350 in PBS, was applied for 1h at RT. Finally, the slides were washed with PBS and covered in Vectashield mounting medium with DAPI (Vector Laboratories).

To calculate the delta ratio between the depletion and the saturation for both sensors, the minimum response (depletion) was obtained during the application of 10 mM pyruvate, and the maximum response (saturation) during the 10 mM lactate stimulation, their difference yielding the delta ratio. To evaluate the effect of different MCT blockers on lactate influx, the positive amplitude peak was measured before and after drug application. The decay time of these responses was calculated as the time required to return to the baseline from the response peak. In both cases, the lactate response and the decay time were first normalized per cell based on the control response and then averaged throughout the experiments. Statistical analysis was performed using GraphPad Prism software (RRID:SCR_002798). The data were first analyzed for normality using the Shapiro–Wilk test. In all cases, the test determined that the data did not conform to a normal distribution, therefore significant differences were established with the Wilcoxon Matched-Pairs Signed Ranks Test. The α value was set to 0.05. Unless otherwise stated, data values are given as mean ± SD. Significance levels as indicated by asterisks were: * p < 0.05; **p < 0.01, *** p < 0.001.

Here, we took advantage of the advantages of the defined and controlled *ex vivo* conditions of organotypic retinal explants to express the FRET nanosensors Laconic (San Martín et al., 2013) and Δ6 (Takanaga et al., 2008). Previously, we had shown that our protocol allowed the culture of mouse retinal explants for up to two weeks without morphological disorganization or major physiological changes (Calbiague et al., 2020). The retinas were cultured at p9, and at p11 and again at p12, the explants were transfected by overnight incubation with one of the nanosensors. Imaging took place after two weeks in culture (Fig. 1A). It is important to note that these sensor have been widely used in different tissues and have been shown to be stable at physiological pH and at room temperature (Mächler et al., 2016; San Martín et al., 2013).

To demonstrate functional nanosensor expression, we measured the ratio between the emissions obtained from monomeric teal fluorescent protein (mTFP)/Venus for Laconic and yellow fluorescent protein/cyan fluorescent protein (YFP/CFP) for Δ6 in the soma (Fig. 1B, D). Then, we saturated the lactate and glucose nanosensors with 10 mM lactate or 10 mM glucose respectively (Fig. 1C, E). To deplete intracellular levels of these metabolites, we took advantage of a property of monocarboxylate transporters (MCTs) called trans-acceleration (Poole and Halestrap, 1993), where an extracellular exposure to a monocarboxylate triggers intracellular substrate efflux. Here, 10 mM pyruvate were applied to deplete intracellular lactate (Fig. 1C). We observed a decrease of the intracellular lactate levels when we changed the solution to 0 mM lactate, indicating that this metabolite flowed in a gradient-dependent way (Fig. 1C). Then, we saturated the lactate and glucose nanosensors with 10 mM lactate or 10 mM glucose respectively (Fig. 1C, E) (Ruminot et al., 2019). Interestingly, when we applied 10 mM glucose, fluorescence levels returned to baseline, suggesting that MCs are working under a saturated intracellular glucose concentration in basal conditions (Fig. 1E). The delta ratio between the depletion and the saturation for Laconic was 17.8 ± 7.2% (Fig. 1C), and for the glucose nanosensor Δ6 38.7 ± 7.6% (Fig. 1E), which is similar to responses reported in other cells and tissues (Ruminot et al., 2019; San Martín et al., 2013). Together, these results demonstrate that different FRET nanosensors can be functionally expressed in retinal explant cultures to study metabolic dynamics on a single-cell level.

The retinal expression patterns of the different MCTs are still unclear. MCTs are encoded by the SLC16A gene family, and act as bidirectional proton-linked carriers of monocarboxylates such as lactate, pyruvate, and ketoacids (Halestrap, 2013). Here, we used the expression of Laconic and complementary immunostaining experiments to determine the expression of MCTs in retinal MCs. Since these transporters work bidirectionally, we applied a pulse of 10 mM lactate to trigger its influx into MCs. The relative amplitude of this response was 13.9 ± 5.2% (Fig. 2A). In mammals, the lactate concentration in the retina fluctuates between 5 and 50 mM depending on the species (Kolko et al., 2016). For this study, we measured the lactate released by retinal explants after 4 days in culture, and obtained a concentration of 16.7 ± 5.7 mM lactate in the culture medium (n = 4), therefore the concentration of lactate used for acute stimulation is unlikely have caused neurotoxic effects.

**Fig 2.**
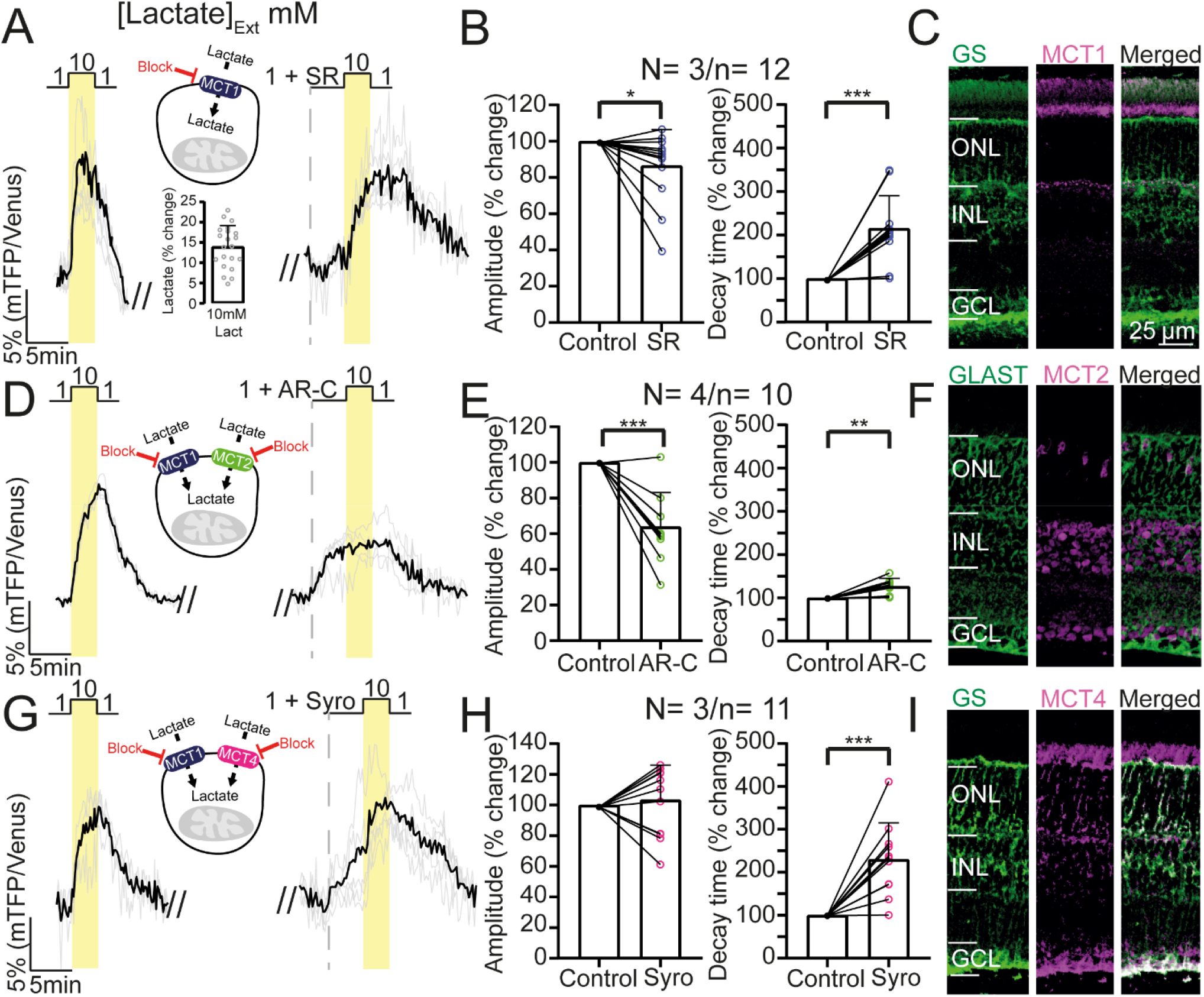
Inhibition of lactate transport disrupts lactate influx in retinal MCs. (A, D, and G, left) Lactate influx evoked after a stimulation with 10 mM lactate. (A, D, and G, right) Lactate influx responses after an incubation with different drugs which target monocarboxylate transporters (MCTs). The light grey traces represent responses of single cells, and the black trace represents their average. (B, E, and H) Statistical analysis of the response amplitude and the decay time of the lactate influx. Paired Student’
ss t-test followed by Wilcoxon’s post hoc. (C, F, and I) Immunofluorescence for different MCTs in the retina showing co-labelling with MC markers. All the images display the same scale bar. Graphs display the mean ± SD. * Indicates p < .05; ** indicates p < .01. *** indicates p < .001. ONL= outer nuclear layer; INL= inner nuclear layer; GCL= ganglion cell layer; MC(s)= Müller cell(s); GLAST= glutamate-aspartate transporter; GS= glutamine synthetase. SR= MCT1 blocker; AR-C= MCT1 and MCT2 blocker; Syro= MCT1 and MCT4 blocker. The number of experiments is represented as: N= number of explants; n= number of cells recorded.

The presence of the specific MCT1 inhibitor SR-13800 0,1 µM (Murray et al., 2005), produced a decrease of 13.4 ± 19.8% in the peak response (Fig. 2B, left). The decay time was even more affected and was extended by 115.6 ± 75.1% (Fig. 2A, B, right). Curiously, the MCT1 immunofluorescence indicated an expression in photoreceptor inner segments and the outer plexiform layer, but failed to show co-labelling with the MC marker glutamine synthetase (GS; Fig. 2C).

When we did the same experiment in the presence of a potent inhibitor of both MCT1 and MCT2, AR-C155858 2 µM (Fig. 2D) (Ovens et al., 2010), the response peak decreased by 36.0 ± 19.2% (Fig. 2D, left), suggesting that lactate influx was mainly mediated by MCT2. We also observed an increase in the decay time by 26.8 ± 18.3% (Fig. 2D, E, right). On the other hand, MCT2 immunostaining displayed strong labelling in the inner nuclear layer (INL), and in some cell bodies in the outer nuclear layer, likely corresponding to cones. Although the co-labelling with the MC-marker glutamate-aspartate transporter (GLAST) did not show a clear colocalization in the INL, GLAST labelling surrounds MCT2 expression in this layer (Fig. 2F).

Finally, when MCT1 and MCT4 were inhibited simultaneously with syrosingopine (4 µM) (Vélez et al., 2021), we did not observe any change in the response amplitudes (Fig. 2G, left). Nonetheless, the decay time had a significant increase of 130.1 ± 85.0% (Fig. 2G, H, right), supporting the hypothesis that MCT1 and MCT4 mediate lactate efflux in MCs. Immunostaining of MCT4 revealed clear co-labelling of MCT4 and the MC marker glutamine synthase (GS), confirming expression of this transporter in MCs (Fig. 2I).

In the present study, we expressed glucose and lactate nanosensors in MCs of organotypic retinal explants to study changes in intracellular lactate concentrations under different conditions. Our observations may provide new insights into retinal metabolism through a live-cell imaging approach. Here, pharmacological data support MCT1, MCT2, and MCT4 isoform expression by MCs, possibly fulfilling different roles in lactate dynamics.

The saturation of Δ6 at basal levels indicates that MCs work under a high intracellular glucose concentration, probably because of the metabolic coupling between these cells and the retinal neurons (Grimes et al., 2021; Toft-Kehler et al., 2018), which rely on MCs for the provision of different metabolites for their metabolic needs. Due to their central role in retinal metabolism and intimate association with blood vessels, MCs likely have high glucose uptake, used in part for the storage of glycogen (Toft-Kehler et al., 2018).

Our data support the expression of different MCT isoforms in MCs. The suggestion of MCT2 expression by MCs is puzzling, since this transporter is known as the neuronal MCT (Pierre et al., 2002), and is indicative of a possible consumption of lactate under certain conditions. In fact, the aforementioned data support MCT2 expression in MCs of mouse retina, in line with previous immunohistochemical data for rats (Gerhart et al., 1999). These results could hence be interpreted as evidence against the ANLS hypothesis in the retina (Kanow et al., 2017; Lindsay et al., 2014). However, the demonstration of MCT4 expression and its role in lactate efflux are indicative of lactate production by MCs (Pellerin and Magistretti, 1994), supporting a complex metabolism in which retinal cells can change between lactate production and consumption, depending on the metabolic demands or the physiological context.

The expression of MCT1 in MCs remains uncertain, even though it was previously detected in rat retinal MCs (Gerhart et al., 1999), as well as very recently through single-cell transcriptomics analysis (Bisbach et al., 2022). However, our immunofluorescence experiments did not confirm MCT1 in MCs, which could be due to different reasons. MCT1 expression might be developmentally regulated during the first weeks of life. As mentioned in methods, the retinal explant cultures were prepared from p9 mice (eyes closed) and maintained in culture for two weeks, while the retinal slices for immunostaining were prepared from p30 mice. This age difference could potentially produce a significant variation in MCT1 expression, given that the peak expression is apparently reached at p21, followed by a reduction in mRNA levels (Clamp et al., 2004). Furthermore, there is the possibility that MCT1 is not expressed in MCs. As observed in the immunolabelling in Fig. 2C and previously reported by others, there is a clear expression in photoreceptor inner segments (Bergersen et al., 1999; Chidlow et al., 2004; Peachey et al., 2018). Therefore, we cannot exclude the possibility that intracellular lactate in MCs might be indirectly affected by photoreceptors metabolically coupled to MCs. It would be relevant to further investigate why MCs might express different types of MCTs, and how the lactate flux through these transporters varies under different visual conditions. Hence, the functional results displayed in Fig. 2 suggest that MCT2, and MCT4 are indeed expressed by MCs, while MCT1 probably is partially expressed at very low levels.

Here, we propose that MCT2 might be in charge of lactate influx and MCT1 and MCT4 of lactate efflux, according to prior suggestions in a high lactate environment (Contreras-Baeza et al., 2019). This suggests that MCs should express the intracellular enzymatic machinery to convert lactate to pyruvate, or pyruvate to lactate. It has been shown that MCs express the lactate dehydrogenase B (LDH-B) isoform (Chinchore et al., 2017). Thus, MCs could convert lactate into pyruvate through LDH-B to fuel mitochondria and produce ATP through oxidative phosphorylation. However, MCs lack the enzymes pyruvate kinase M2 (PKM2) and lactate dehydrogenase A (LDH-A) required for lactate production (Chen et al., 2022; Chinchore et al., 2017; Lindsay et al., 2014; Rajala et al., 2018). It is possible that the expression of pyruvate kinase M1 (PKM1) and LDH-B are sufficient to sustain lactate synthesis in MCs and release it through MCT1 and MCT4 when metabolically required. Certainly, more direct evidence is required to test this idea in the future.

It is important to indicate the limitations of our study. All measurements were obtained from cell bodies, whereas the long cellular projections of MCs might be metabolically isolated and display different metabolite dynamics. Furthermore, the experimental conditions used here might will not accurately reflect the physiological conditions of *in vivo* retina. While retinas were light-adapted, photoreceptors can show variations in their activity after prolonged time in culture depending on the status of the RPE (Alarautalahti et al., 2019; Tolone et al., 2021).

In summary, our physiological and immunohistochemical results support expression and function of MCT2 and MCT4 in MCs, while the expression of MCT1 in these cells, although suggested by pharmacological data, is not confirmed by immunohistochemisty. The proof of concept of glucose and lactate nanosensor expression and function in organotypic retinal explants opens a wide range of experiments which will contribute to further understand the complex retinal metabolism under physiological and pathological conditions.

## 2 Funding

This study was supported by FONDECYT grant No. 1210790 and ANID PhD scholarship No. 21180443 to VCG.

## 3 Author Contributions

VCG and OS contributed to the conception and design of the study. VCG and YC contributed to data acquisition, analysis, and figures preparation. VCG, YC, BC, FPD and OS contributed to article writing and approved the submitted version.

## 4 Acknowledgments

We wish to thank Dr. Ivan Ruminot, Dr. Felipe Baeza-Lehnert, and Dr. Felipe Barros from Centros de Estudios Científicos (CECs) in Valdivia (Chile) for sharing its expertise in FRET measurements, and to Felipe Tapia from Universidad de Valparaiso for helping in the data analysis.

## 5 Declaration of competing interest

The authors declare no competing interests

## 6 Ethics statements

The animal study was reviewed and approved by Comité institucional para el cuidado y uso de animales de laboratorio (CICUAL), Universidad de Valparaiso

## Notes

### Competing Interest Statement

The authors have declared no competing interest.

## References

Alarautalahti, V., Ragauskas, S., Hakkarainen, J.J., Uusitalo-Järvinen, H., Uusitalo, H., Hyttinen, J., Kalesnykas, G., Nymark, S., 2019. Viability of mouse retinal explant cultures assessed by preservation of functionality and morphology. Investig. Ophthalmol. Vis. Sci. 60, 1914–1927. https://doi.org/10.1167/iovs.18-25156

Ames, A., Li, Y.Y., Heher, E.C., Kimble, C.R., 1992. Energy metabolism of rabbit retina as related to function: High cost of Na+ transport. J. Neurosci. 12, 840–853. https://doi.org/10.1523/jneurosci.12-03-00840.1992

Barros, L.F., Baeza-Lehnert, F., Valdebenito, R., Ceballo, S., Alegría, K., 2014. Fluorescent Nanosensor Based Flux Analysis: Overview and the Example of Glucose., in: In Brain Energy. humana press, New york, pp. 145–159.

Belhadj, S., Tolone, A., Christensen, G., Das, S., Chen, Y., Paquet-Durand, F., 2020. Long-term, serum-free cultivation of organotypic mouse retina explants with intact retinal pigment epithelium. J. Vis. Exp. 2020, 1–13. https://doi.org/10.3791/61868

Bergersen, L., Jo, E., Veruki, M.L., Nagelhus, E.A., 1999. Cellular and subcellular expression of monocarboxylate transporters in the pigment epithelium and retina of the rat. Neuroscience 90, 319–331.

Bisbach, C.M., Hass, D.T., Thomas, E.D., Cherry, T.J., Hurley, J.B., Ed, T., Tj, C., Jb, H., 2022. Monocarboxylate Transporter 1 (MCT1) mediates succinate export in the retina. Invest. Ophthalmol. Vis. Sci. 63, 1–1.

Calbiague, V.M., Vielma, A.H., Cadiz, B., Paquet-Durand, F., Schmachtenberg, O., 2020. Physiological assessment of high glucose neurotoxicity in mouse and rat retinal explants. J. Comp. Neurol. 528, 989–1002. https://doi.org/10.1002/cne.24805

Chen, Y., Zizmare, L., Calbiague, V., Yu, S., Herberg, F.W., 2022. Retinal energy metabolismL: Photoreceptors switch between Cori, Cahill, and mini-Krebs cycles to uncouple glycolysis from mitochondrial respiration. bioRxiv.

Chidlow, G., Wood, J.P.M., Graham, M., Osborne, N.N., Wood, J.P.M., Graham, M., 2004. Expression of monocarboxylate transporters in rat ocular tissues. Am. J. Physiol. Physiol. 288, 416–428. https://doi.org/10.1152/ajpcell.00037.2004.

Chinchore, Y., Begaj, T., Wu, D., Drokhlyansky, E., Cepko, C.L., 2017. Glycolytic reliance promotes anabolism in photoreceptors. Elife 6, 1–22. https://doi.org/10.7554/eLife.25946

Clamp, M.F., Ochrietor, J.D., Moroz, T.P., Linser, P.J., 2004. Developmental analyses of 5A11/Basigin, 5A11/Basigin-2 and their putative binding partner MCT1 in the mouse eye. Exp. Eye Res. 78, 777–789. https://doi.org/10.1016/j.exer.2003.12.004

Contreras-Baeza, Y., Sandoval, P.Y., Alarcón, R., Galaz, A., Cortés-Molina, F., Alegriá, K., Baeza-Lehnert, F., Arce-Molina, R., Guequén, A., Flores, C.A., Martín, A.S., Barros, L.F., 2019. Monocarboxylate transporter 4 (MCT4) is a high affinity transporter capable of exporting lactate in high-lactate microenvironments. J. Biol. Chem. 294, 20135–20147. https://doi.org/10.1074/jbc.RA119.009093

Country, M.W., 2017. Retinal metabolismL: A comparative look at energetics in the retina. Brain Res. 1672, 50–57. https://doi.org/10.1016/j.brainres.2017.07.025

Gerhart, D.Z., Leino, R.L., Drewes, L.R., 1999. Distribution of monocarboxylate transporters MCT1 and MCT2 in rat retina. Neuroscience 92, 367–375.

Grimes, W.N., Aytürk, D.G., Hoon, M., Yoshimatsu, T., Gamlin, C., Carrera, D., Nath, A.x, Nadal-Nicolás, F.M., Ahlquist, R.M., Sabnis, A., Berson, D.M., Diamond, J.S., Wong, R.O., Cepko, C., Rieke, F., 2021. A high-density narrow-field inhibitory retinal interneuron with direct coupling to Müller glia. J. Neurosci. 41, 6018–6037. https://doi.org/10.1523/JNEUROSCI.0199-20.2021

Halestrap, A.P., 2013. Monocarboxylic Acid Transport. Compr. Physiol. 3, 1611–1643. https://doi.org/10.1002/cphy.c130008

Kanow, M.A., Giarmarco, M.M., Jankowski, C.S.R., Tsantilas, K., Engel, A.L., Du, J., Linton, J.D., Farnsworth, C.C., Sloat, S.R., Rountree, A., Sweet, I.R., Lindsay, K.J., Parker, E.D., Brockerhoff, S.E., Sadilek, M., Chao, J.R., Hurley, J.B., 2017. Biochemical adaptations of the retina and retinal pigment epithelium support a metabolic ecosystem in the vertebrate eye. Elife 6, 1–25. https://doi.org/10.7554/eLife.28899

Kolko, M., Vosborg, F., Henriksen, U.L., Hasan-Olive, M.M., Diget, E.H., Vohra, R., Gurubaran, I.R.S., Gjedde, A., Mariga, S.T., Skytt, D.M., Utheim, T.P., Storm-Mathisen, J., Bergersen, L.H., 2016. Lactate Transport and Receptor Actions in Retina: Potential Roles in Retinal Function and Disease. Neurochem. Res. 41, 1229–1236. https://doi.org/10.1007/s11064-015-1792-x

Lindsay, K.J., Du, J., Sloat, S.R., Contreras, L., Linton, J.D., Turner, S.J., Sadilek, M., Satrústegui, J., Hurley, J.B., 2014. distributions reveal key metabolic links between neurons and glia in retina. Proc. Natl. Acad. Sci. 111, 15579–15584. https://doi.org/10.1073/pnas.1412441111

Mächler, P., Wyss, M.T., Elsayed, M., Stobart, J., Gutierrez, R., Von Faber-Castell, A., Kaelin, V., Zuend, M., San Martín, A., Romero-Gómez, I., Baeza-Lehnert, F., Lengacher, S., Schneider, B.L., Aebischer, P., Magistretti, P.J., Barros, L.F., Weber, B., 2016. In Vivo Evidence for a Lactate Gradient from Astrocytes to Neurons. Cell Metab. 23, 94–102. https://doi.org/10.1016/j.cmet.2015.10.010

Murray, C.M., Hutchinson, R., Bantick, J.R., Belfield, G.P., Benjamin, A.D., Brazma, D., Bundick, R. V., Cook, I.D., Craggs, R.I., Edwards, S., Evans, L.R., Harrison, R., Holness, E., Jackson, A.P., Jackson, C.G., Kingston, L.P., Perry, M.W.D., Ross, A.R.J., Rugman, P.A., Sidhu, S.S., Sullivan, M., Taylor-Fishwick, D.A., Walker, P.C., Whitehead, Y.M., Wilkinson, D.J., Wright, A., Donald, D.K., 2005. Monocarboxylate Transporter Mctl is a Target for Immunosuppression. Nat. Chem. Biol. 1, 371–376. https://doi.org/10.1038/nchembio744

Ovens, M.J., Davies, A.J., Wilson, M.C., Murray, C.M., Halestrap, A.P., 2010. AR-C155858 is a potent inhibitor of monocarboxylate transporters MCT1 and MCT2 that binds to an intracellular site involving transmembrane helices 7-10. Biochem. J. 425, 523–530. https://doi.org/10.1042/BJ20091515

Peachey, N.S., Yu, M., Han, J.Y.S., Lengacher, S., Magistretti, P.J., Pellerin, L., Philp, N.J., 2018. Impact of MCT1 haploinsufficiency on the mouse retina. Adv. Exp. Med. Biol. 1074, 375–380. https://doi.org/10.1007/978-3-319-75402-4_46

Pellerin, L., Magistretti, P.J., 1994. Glutamate uptake into astrocytes stimulates aerobic glycolysis: A mechanism coupling neuronal activity to glucose utilization. Proc. Natl. Acad. Sci. U. S. A. 91, 10625–10629. https://doi.org/10.1073/pnas.91.22.10625

Pierre, K., Magistretti, P.J., Pellerin, L., 2002. MCT2 is a major neuronal monocarboxylate transporter in the adult mouse brain. J. Cereb. Blood Flow Metab. 22, 586–595. https://doi.org/10.1097/00004647-200205000-00010

Poitry-Yamate, C.L., Poitry, S., Tsacopoulos, M., 1995. Lactate Released by Miiller Glial Cells Is Metabolized Photoreceptors from Mammalian Retina. J. Neurosci. 15, 5179–5191.

Poole, R.C., Halestrap, A.P., 1993. Transport of lactate and other monocarboxylates across mammalian plasma membranes. Am. J. Physiol. - Cell Physiol. 264, 761–782. https://doi.org/10.1152/ajpcell.1993.264.4.c761

Rajala, A., Wang, Y., Brush, R.S., Tsantilas, K., Jankowski, C.S.R., Lindsay, K.J., Linton, J.D., Hurley, J.B., Anderson, R.E., Rajala, R.V.S., 2018. Pyruvate kinase M2 regulates photoreceptor structure, function, and viability article. Cell Death Dis. 9. https://doi.org/10.1038/s41419-018-0296-4

Ruminot, I., Schmälzle, J., Leyton, B., Barros, L.F., Deitmer, J.W., 2019. Tight coupling of astrocyte energy metabolism to synaptic activity revealed by genetically encoded FRET nanosensors in hippocampal tissue. J. Cereb. Blood Flow Metab. 39, 513–523. https://doi.org/10.1177/0271678X17737012

San Martín, A., Ceballo, S., Ruminot, I., Lerchundi, R., Frommer, W.B., Barros, L.F., 2013. A Genetically Encoded FRET Lactate Sensor and Its Use To Detect the Warburg Effect in Single Cancer Cells. PLoS One 8. https://doi.org/10.1371/journal.pone.0057712

Takanaga, H., Chaudhuri, B., Frommer, W.B., 2008. GLUT1 and GLUT9 as major contributors to glucose influx in HepG2 cells identified by a high sensitivity intramolecular FRET glucose sensor. Biochim. Biophys. Acta (BBA)-Biomembranes 1778, 1091–1099. https://doi.org/10.1201/b18990-115

Toft-Kehler, A.K., Skytt, D.M., Kolko, M., 2018. A Perspective on the Müller Cell-Neuron Metabolic Partnership in the Inner Retina. Mol. Neurobiol. 55, 5353–5361. https://doi.org/10.1007/s12035-017-0760-7

Tolone, A., Haq, W., Fachinger, A., Rentsch, A., Herberg, F.W., Schwede, F., Paquet-Durand, F., 2021. Retinal degeneration: Multilevel protection of photoreceptor and ganglion cell viability and function with the novel PKG inhibitor CN238. bioRxiv.

Tsacopoulos, M., Evêquoz-Mercier, V., Perrottet, P., Buchner, E., 1988. Honeybee retinal glial cells transform glucose and supply the neurons with metabolic substrate. Proc. Natl. Acad. Sci. U. S. A. 85, 8727–8731. https://doi.org/10.1073/pnas.85.22.8727

Valdés, J., Trachsel-Moncho, L., Sahaboglu, A., Trifunović, D., Miranda, M., Ueffing, M., Paquet-Durand, F., Schmachtenberg, O., 2016. Organotypic retinal explant cultures as in vitro alternative for diabetic retinopathy studies. ALTEX 33, 459–464. https://doi.org/10.14573/altex.1603111

Vélez, L., Velasquez, Z., Silva, L.M.R., Gärtner, U., Failing, K., Daugschies, A., Mazurek, S., Hermosilla, C., Taubert, A., 2021. Metabolic Signatures of Cryptosporidium parvum -Infected HCT-8 Cells and Impact of Selected Metabolic Inhibitors on. Biology (Basel). 10, 60.

Wang, L., Tornquist, P., Billl, A., 1997. Glucose metabolism in pig outer retina in light and darkness. Acta Physiol. Scand. 160, 75–81.

Warburg, O., 1924. The metabolism of carcinoma cells. J. Cancer Res. 9, 148–163.

